# SIRT2 Mediates Integrated Stress Response by Deacetylating and Stabilizing 4EBP1 to Suppress Translation

**DOI:** 10.1101/2025.03.31.646365

**Authors:** Yanlin Zi, Maio Wang, Dan Hou, Richard A. Cerione, Hening Lin

**Author notes:** The authors contributed equally as first authors (Yanlin Zi, Maio Wang). Disclosures: HL is an inventor on Cornell University’s patent applications on SIRT2 inhibitors.

## Abstract

The ability to alleviate nutrient stress, such as amino acid limitation, is crucial for cell survival. The mTORC1 complex and integrated stress response (ISR) are mechanisms that sense the availability of amino acids and regulate ribosomal protein synthesis. Here we have discovered a new SIRT2-mediated pathway, downstream of ISR, that senses the limitation of amino acids to regulate translation. Under amino acid deprivation, SIRT2 protein level is upregulated translationally by its upstream open reading frame (uORF). SIRT2 in turn suppresses global protein translation, which helps cells to survive amino acid limitation. Mechanistically, we identified eukaryotic translation initiation factor 4E (eIF4E) binding protein 1 (4EBP1), which negatively regulates translation, as a substrate of SIRT2. SIRT2 deacetylates 4EBP1 at Lys69 and stabilizes 4EBP1 by protecting it from proteasomal degradation. Our study reveals a novel role for SIRT2 in regulating protein translation and a new regulatory mechanism of 4EBP1 in cells. Our study provides a better understanding of the intricate regulation of translation and may explain the known non-oncogene addiction role of SIRT2 in cancer cells.

## Introduction

Stress response pathways are crucial for cell survival, and they are often heavily relied on by cancer cells, which are constantly under various stress conditions^1^. Cancer cells must adapt to nutrient and oxygen limitation, proteotoxic stress, and DNA damage, all caused by their high demand for proliferation. Therefore, stress response pathways have become part of cancer cells’ non-oncogene addiction^2^. By targeting proteins involved in this form of non-oncogene addiction, such as stress response proteins, tumors can be selectively targeted while sparing healthy cells. If the intricacies of the stress response pathways can be better understood, it would aid the design and development of drugs targeting diseases including cancer.

The integrated stress response (ISR) pathway is an elaborated signaling pathway that enables cells to respond and handle stresses from amino acid deprivation, viral infection, heme deprivation, and ER stress^3^. When cells sense one of these four types of stresses, an ISR regulator GCN2 for amino acid stress, PKR for viral infection, HRI for heme stress, and PERK for ER stress are activated and trigger the phosphorylation of eukaryotic translation initiation factor 2α (eIF2α) on Ser51. This phosphorylation of eIF2α attenuates global cap-dependent mRNA translation but initiates the preferential translation of certain mRNAs that contain upstream open reading frames (uORF), such as activating transcription factor 4 (ATF4), as a mechanism for promoting cell survival and recovery^4^.

Like other stress adaptation pathways, defects in ISR have also been shown to suppress cancer and other diseases. Inhibition of eIF2α Ser51 phosphorylation was reported to trigger cytotoxicity in aggressive metastatic prostate cancer^5^. ATF4, one of the few well-studied uORF-regulated proteins in ISR, was reported to be overexpressed in solid tumors and critical for maintaining metabolic homeostasis in tumor cells^6^. This underlies the importance of identifying more proteins directly regulated by ISR and to better understand the stress responses of cancer cells.

mTORC1 regulates a number of critically important cellular processes. It affects cell growth by regulating mRNA translation, metabolism, and protein turnover^7^. Like the ISR pathway, mTORC1 also senses growth factors and specific amino acids, such as leucine and arginine^8,9^ as signaling cues to promote cell growth and survival. Thus, when arginine or leucine levels are low, mTORC1 activity decreases. mTORC1 regulates protein translation through its phosphorylation of ribosomal S6 kinase (S6K), which induces translation^10,11^, and eukaryotic translation initiation factor 4E (eIF4E)-binding protein 1 (4EBP1), which negates the inhibitory effects it would otherwise have on translation^12,13^.

Sirtuins are NAD^+^ dependent deacetylases. Within the sirtuin family, there are seven members, SIRT1-7, in humans^14^. Although they are highly conserved, each sirtuin has its own localization and cellular function^15^. In this study, we mainly focus on SIRT2, which was discovered over two decades ago^16^, but whose complete function is still not fully understood. In addition to being capable of exhibiting strong deacetylase activity, SIRT2 has also been reported to have defatty-acylase activity^17–21^. It has been shown to have important roles in regulating gene expression^22,23^, signal transduction^24–26^, metabolic disorders^27,28^, and various diseases including cancer^29^, diabetes^30^, inflammatory bowel disease (IBD)^31^ and neurodegenerative diseases^32^. Eukaryotic translation initiation factor 5A (eIF5A) has been reported to be a deacetylation substrate of SIRT2, but whether and how SIRT2 regulates translation is not well known^33^.

In this study, we demonstrate that SIRT2 is translationally upregulated under amino acid stress via ISR and that SIRT2 negatively regulates global protein translation. Moreover, we discovered that 4EBP1 is a substrate of SIRT2. When 4EBP1 undergoes SIRT2 catalyzes deacetylation at lysine 69 (K69), it is then protected from proteasomal degradation, which contributes to translational suppression.

## Results

### SIRT2 protein level is upregulated under amino acid limitation

We first discovered that SIRT2 protein levels were increased under conditions of amino acid depletion. In multiple cell lines, including cancer cells and non-cancer cells such as MEFs and HEK293T cells, SIRT2 was consistently upregulated under lysine (K) and arginine (R), or leucine, lysine and arginine (LKR) deprivation (Figure 1A and 1B). Next, we tested more amino acid depletion conditions. SIRT2 upregulation was observed under all conditions examined, including deprivation of branch chain amino acids (leucine), charged amino acids (lysine or arginine), and non-essential amino acids (serine and glycine) for 12 or 24 hours (Figure 1C). Taken together, the data suggests that increased SIRT2 protein levels as a response to amino acids deprivation is a general phenomenon.

**Figure 1.**
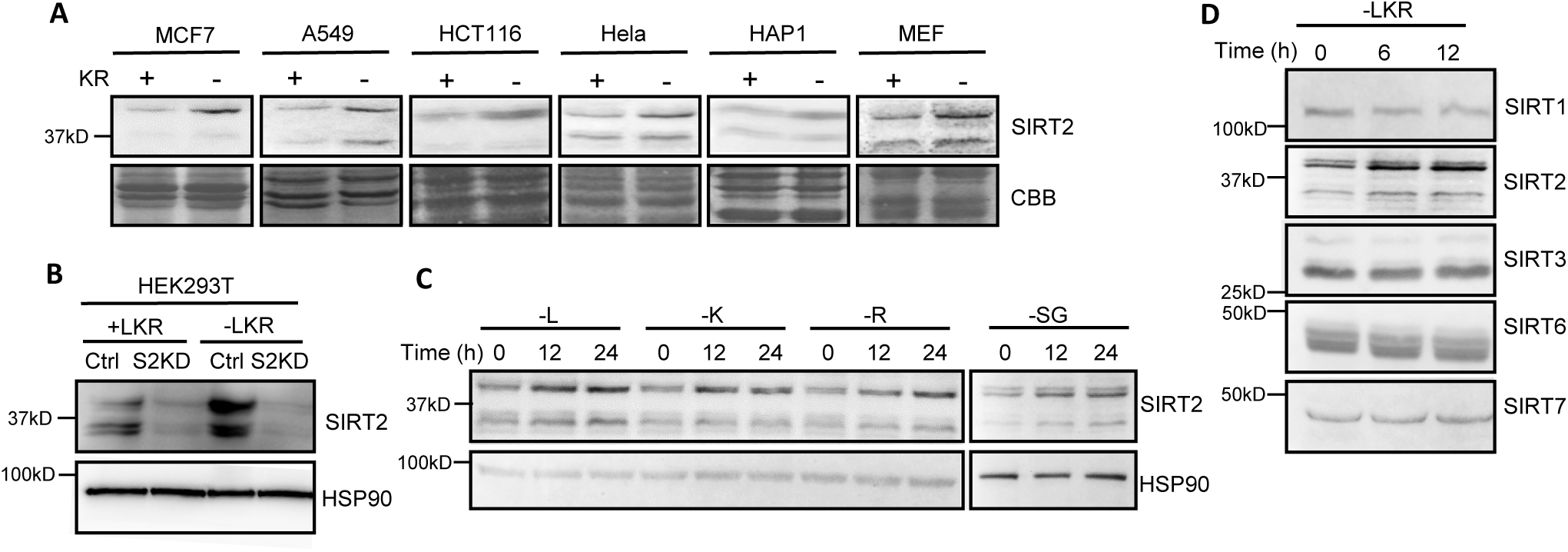
SIRT2 is upregulated under amino acid limitation. **(A)** SIRT2 level is upregulated in multiple cell lines when starved with lysine (K) and arginine (R) for 24 hrs. SIRT2 levels in MCF7, A549, HCT116, Hela, HAP1, and MEF cells with or without lysine and arginine were measure by western blot. CBB: Coomassie Brilliant Blue staining of total proteins. **(B)** SIRT2 level is upregulated in HEK293T cells when subjected to leucin, lysine, arginine (LKR) starvation for 24 hrs. Western blot was used to determine SIRT2 levels. SIRT2 knockdown HEK293T cells were negative controls. **(C)** Immunoblots for SIRT2 levels in MCF cells starved of leucine (L), lysine (K), arginine (R), and serine (S) and glycine (G) for 12 and 24 hrs. Protein levels were evaluated using western blot. **(D)** Only SIRT2 level increases under lysine (L), leucine (K), and arginine (R) depletion. SIRT1/3/6/7 protein level is not affected by LKR depletion. MCF7 cells were starved of LKR for 6 and 12 hrs. Western blot was used to determine the sirtuin protein levels.

We next examined the protein levels of several other sirtuins in response to amino acid deprivation. Among the sirtuins we examined (SIRT1/2/3/6/7), SIRT2 was the only sirtuin that was elevated under leucine, lysine and arginine deprivation (Figure 1D), suggesting that SIRT2 has a unique role when cells are under amino acid deprivation.

### Upregulation of SIRT2 translation is controlled translationally by its uORFs

Amino acid deprivation is one of the four types of stresses that activate the integrated stress response (ISR), and the best understood downstream response of ISR is the upregulation of ATF4 via translational control^35^. When cells detect amino acid deprivation, ATF4 activates the transcription of its target genes, including itself, to assist cells in overcoming stress. Given that SIRT2 protein levels increased in response to amino acid deprivation, which is a stress that activates ISR, we then investigated whether this upregulation occurred transcriptionally or translationally. If the upregulation of SIRT2 occurs at a transcription level, then it would imply that SIRT2 is an ATF4 target gene. If the upregulation occurs at a translation level, then it is possible that SIRT2 translation is regulated similarly to the regulation of ATF4 translation when cells are under amino acid stress.

To answer this question, we first evaluated *SIRT2* transcript abundance following amino acid deprivation using real time (RT)-PCR. A549 cells were starved of leucine, lysine, and arginine for 1, 6, 12, and 24 hours before collection. As expected, both the protein and relative mRNA levels of ATF4 increased during the time course. However, for SIRT2, only the protein level of SIRT2 increased upon the deprivation of amino acids LKR (Figure 2A). Similarly, when deprived of glutamine (Q), lysine and arginine (KR), or all three combined (QKR), the relative *SIRT2* mRNA levels remained the same compared to the sample without any amino acid deprivation, even though SIRT2 protein levels had increased as observed from the western blot result (Figure 2B). These results showed that the upregulation of SIRT2 protein level is not due to increased transcription.

**Figure 2.**
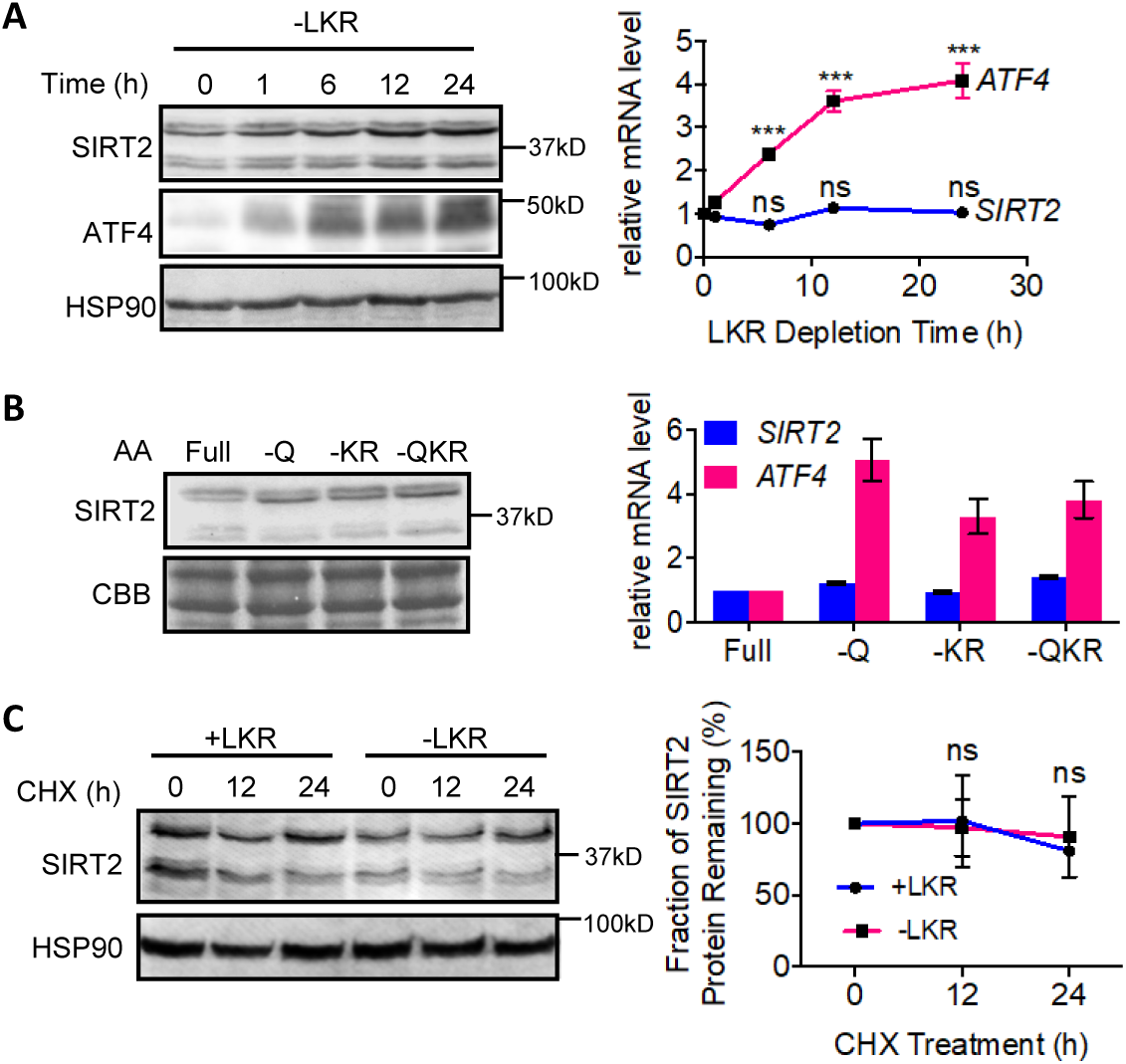
Increased SIRT2 protein level is not due to changes in transcription or stability. **(A)** SIRT2’s relative mRNA levels remain the same under leucine, lysine, and arginine (LKR) depletion. Western blots (left) shows that SIRT2 protein level in A549 cells increased under LKR depletion for various time points. Real-time PCR analysis (right) shows that the relative SIRT2 mRNA level remained the same. **(B)** SIRT2 mRNA levels remain the same under different combinations of amino acids depletion. A549 cells were treated with starvation of glutamine (Q), lysine and arginine (KR), or a combination of all three (QKR) for 24 hrs. Western blots (left) were used to detect SIRT2 protein level, and real-time PCR analysis (right) was used to show relative mRNA levels. **(C)** LKR depletion does not affect the stability of SIRT2. Cycloheximide (CHX) chase experiment was utilized. A549 cells were treated with 50 μg/ml CHX for 0, 12, or 24 hrs. For the western blot quantification (right), SIRT2 protein level from the sample without CHX is set to 1. Data are represented as mean ± SEM. ***p<0.001; ns, not significant. Representative images from 2-3 independent experiments are shown.

We next examined SIRT2 protein stability to verify whether the increased protein level is due to increased translation or stability. To test protein stability, A549 cells were treated with 50 μg/ml of protein synthesis inhibitor cycloheximide (CHX) for 0, 12, or 24 hours, and the SIRT2 protein levels were monitored. There was no significant difference in SIRT2 stability between the LKR starved and the amino acid rich conditions (Figure 2C). Therefore, changes in SIRT2 protein stability did not account for the increased SIRT2 protein level under amino acid depletion.

After ruling out the regulation of transcription and SIRT2 protein stability, it became highly possible that increased SIRT2 translation is responsible for the upregulated SIRT2 protein level upon amino acid deprivation. Under conditions of ISR, eIF2α is phosphorylated on Ser51 and becomes inactive. This inactivation leads to a decreased concentration of the ternary complex which allows scanning ribosomes to skip the inhibitory uORF of specific mRNA, such as Atf4, and results in increased translation of those mRNA transcripts^36^. Therefore, we hypothesized that the translation of SIRT2 is regulated similarly to that of ATF4. To test our hypothesis, we put a luciferase reporter under control of SIRT2 5’ untranslated region (5’UTR) to evaluate the translation efficiency^37^, as the luciferase assay is a well-established tool in the study of integrated stress response mediated translation change^38,39^.

The 5’UTR from the human SIRT2 variant 1 (NM_012237.3) was inserted into the pGL3 vector driven by the SV40 promoter. The start codon of the SIRT2 main open reading frame (mORF) was fused in frame with firefly luciferase. For an internal control, we used a renilla luciferase reporter construct which was under the same promoter. The two constructs were co-transfected into Hela cells and the SIRT2 5’UTR regulated translation efficiency was measured by the ratio of firefly luciferase activity over renilla luciferase activity (Figure 3A and 3B). In the presence of SIRT2 5’UTR, the translation efficiency increased after amino acid depletion or ER stress induction in a time-dependent manner, while the transcription level remains similar (Figure 3A and 3C). This result directly confirmed that upregulation of SIRT2 is through translational regulation, likely via the uORFs in the 5’UTR region of the SIRT2 mRNA.

**Figure 3.**
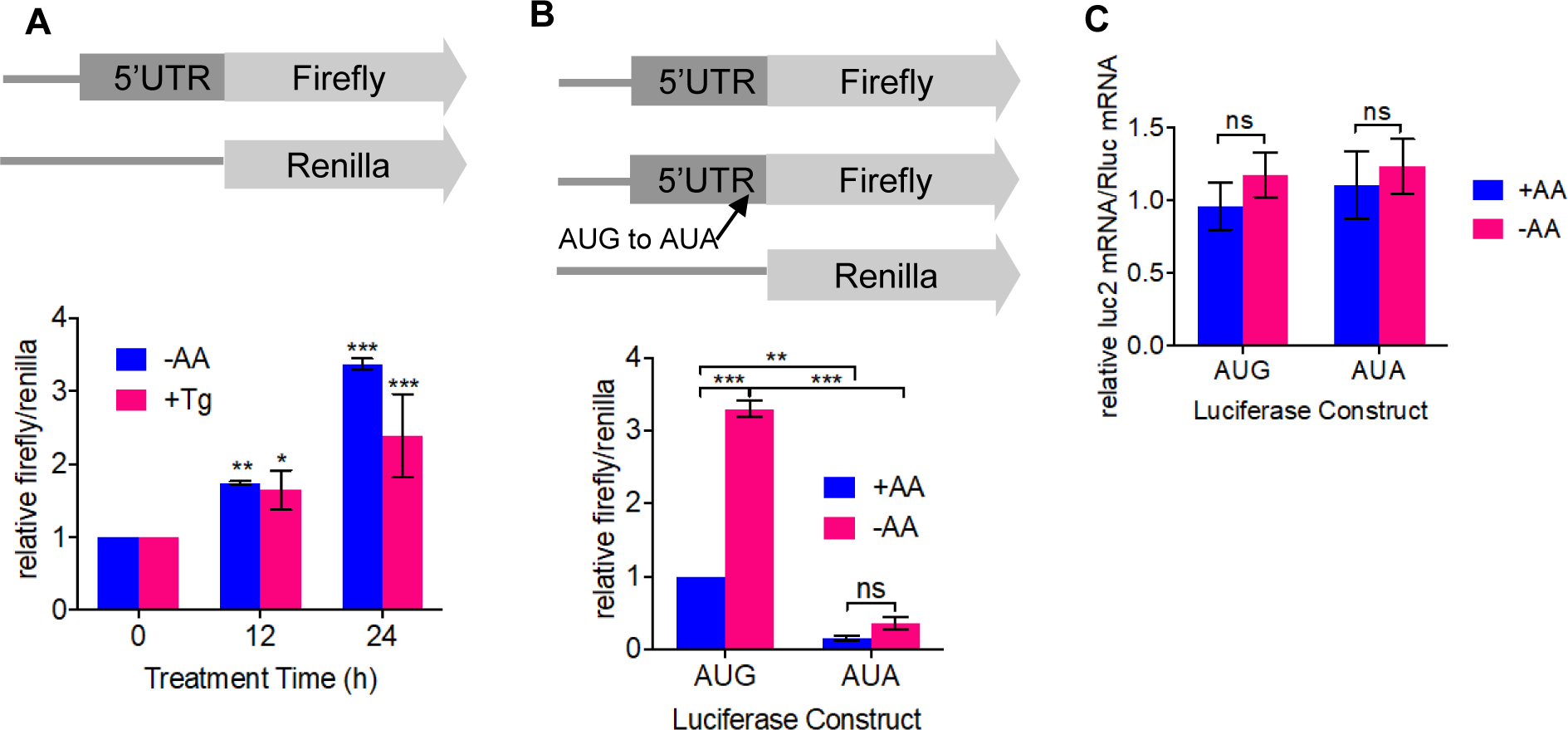
Upregulation of SIRT2 translation is mediated by its uORFs. **(A)** Top: luciferase constructs, 5’ UTR of Sirt2 is inserted between HindIII and NcoI restriction sites of pGL3. Bottom: the ratio of firefly luciferase activity over renilla luciferase activity in Hela cells with LKR depletion or 1 mM thapsigargin (Tg) treatment. **(B)** Top: luciferase constructs with AUG at position 133 mutated to AUA. Bottom: the ratio of firefly luciferase activity over renilla luciferase activity in Hela cells starved of LKR. (C) mRNA level of luciferase reporter assay. Statistical evaluation was done using a two-way ANOVA. Data are represented as mean ±SEM. ***p<0.001.

The most well studied case of the uORF regulating translation is yeast Gcn4 and both inhibitory and activating uORFs have been established^40^. When the inhibitory uORF (uORF4) is lost, ribosome can scan to the main ORF, resulting in a constitutively induced protein translation.

When the activating uORF (uORF1) is lost, the ribosome dissociates more readily from mRNA, resulting in a non-inducible Gcn4 translation. We predicted potential upstream open reading frames in the 5’UTR of SIRT2 mRNA based on previous reports of ribosome profiling analysis^41^ and performed mutagenesis to remove the corresponding start or stop codons. We identified one uORF (133-234) (Figure S1) that when mutated to the non-Kozak sequence, resulted in dramatically decreased translation (Figure 3B). This suggests that the uORF (133-234) in SIRT2 serves a stimulatory role similar to uORF1 in Gcn4. Thus, the translational upregulation of SIRT2 under amino acid depletion is regulated by the uORF in the 5’ UTR.

### Knocking down SIRT2 promotes protein translation and downregulates 4EBP1 protein level

The next question is why is SIRT2 upregulated under conditions of amino acid stress. What functions of SIRT2 are needed when cells are exposed to amino acid stress? Since protein translation is important for cell survival and often regulated by environmental factors such as amino acid abundance^42^, we proceeded to verify if SIRT2 has any effect on protein translation. A previously established puromycin labeling method to probe global protein synthesis was used^43^. Cells were incubated with 10 μg/ml puromycin for 10 minutes before collection, with the immunoblotting signal of puromycin then being directly proportional to the amount of global protein synthesis. We observed that when SIRT2 was knocked out in MEF cells, there was a significant increase in puromycin labeling, indicating that depletion of SIRT2 can increase translation (Figure 4A). Similar results were also observed in HEK293T cells. Knocking down SIRT2 expression in HEK293T cells gave rise to increased puromycin labeling (Figure 4B). We further confirmed this finding in experiments where SIRT2 was overexpressed or its enzymatic activity inhibited. Inhibition of SIRT2 using the small molecule inhibitor TM^44^ increased translation, while overexpression of SIRT2 decreased translation (Figure S2). Thus, SIRT2 has an inhibitory effect on global translation. This could explain why SIRT2 levels are upregulated under amino acid deprivation. When amino acids are limited, cells need to decrease translation. Therefore, when amino acids are scarce, SIRT2 protein level increases to down-regulate translation.

**Figure 4.**
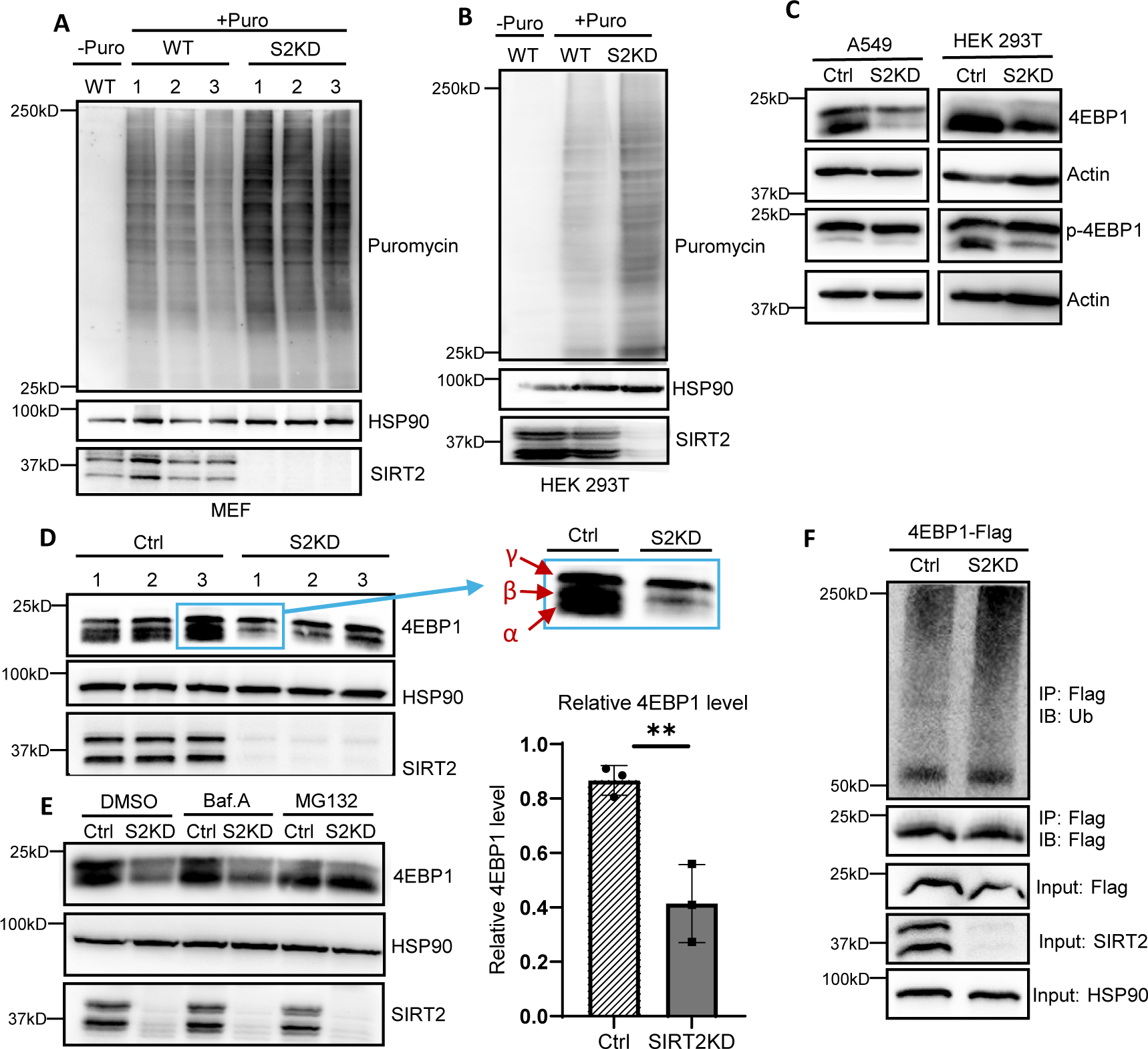
Knocking down SIRT2 promotes protein translation and downregulates 4EBP1 through proteasome degradation. (A-B) SIRT2 knockdown MEF cells **(A)** and HEK293T cells **(B)** have upregulated protein translation. Wildtype (WT) and SIRT2 knockdown (S2KD) cells were treated with10 μg/ml puromycin for 10 minutes before collection. Protein translation level were measured using western blot. **(C)** SIRT2 knockdown A549 and HEK293T cells have decreased 4EBP1 level. Protein levels were measured using western blot. **(D)** SIRT2 knockdown HEK293T cells have significantly reduced α isoform of 4EBP1. Endogenous protein levels in control and SIRT2 knockdown HEK293T cells were evaluated using western blot. Relative 4EBP1 levels to control protein levels were quantified and analyzed. Error bars represent ±SD for experiments performed in triplicate. ** indicating p<0.01. **(E)** MG132 treatment could rescue 4EBP1 level in SIRT2 knockdown HEK293T cells. Control and SIRT2 knockdown HEK293T cells were treated with DMSO (control), 100 nM Bafilomycin A1, or 40 μM MG132 for 5 hrs before collection. Protein levels were evaluated using western blot. **(F)** Knockdown of SIRT2 increases 4EBP1 ubiquitination. Flag-tagged 4EBP1 was transfected into control and SIRT2 knockdown HEK293T cells and was pulled down by Flag beads. Cells were treated with 40 μM MG132 for 6 hours before collection. Ubiquitination level was analyzed using western blot.

To further study SIRT2’s role in translation, we set out to identify a specific target that is important for protein translation and is affected by SIRT2 directly. Translation can be regulated through S6K, whose phosphorylation by mTORC1 will stimulate protein translation^45,46^, and 4EBP1, which exerts an inhibitory effect on protein translation when it is not phosphorylated by binding to the eukaryotic translation initiation factor 4E (eIF4E)^13,47^. We observed that when SIRT2 was knocked down in A549 and HEK293T cells, while phosphorylated 4EBP1 levels remained unchanged, endogenous 4EBP1 protein level was significantly decreased, which could increase global translation (Figure 4C). Since SIRT2 has no significant effect on endogenous S6K expression (Figure S3), we decided to further examine the relationship between SIRT2 and 4EBP1.

### Knocking down SIRT2 promotes the proteasomal degradation of the *α* isoform of 4EBP1

Based on western blot analysis, three 4EBP1 isoforms can be identified: α, β, and γ. The α isoform is the least phosphorylated and migrates the furthest when examined by SDS-PAGE and Western blotting. The β isoform is phosphorylated at intermediate level, and γ isoform is hyperphosphorylated. It is the α isoform of 4EBP1 that binds to eIF4E to inhibit protein translation^48–50^. When using a higher percentage gel to obtain better separation for small proteins, we observed that when SIRT2 was knocked down in HEK293T cells, the total 4EBP1 level decreased by roughly 50% compared to the control HEK293T cells (Figure 4D). The primary protein level difference was with the α isoform. When SIRT2 was knocked down, virtually all of the 4EBP1 α isoform was eliminated while the γ isoform was largely unaffected. This finding is consistent with the observation that knocking down SIRT2 promotes global protein translation, given that the 4EBP1 α isoform inhibits translation.

We then examined how SIRT2 downregulated 4EBP1. Given that SIRT2 is reported to affect the degradation of its targets^51,52^, we hypothesized that the downregulation of 4EBP1 could also be due to increased degradation. To test this hypothesis, HEK293T cells were treated with DMSO (control), 100 nM Bafilomycin A1 for lysosomal inhibition, or 40 μM MG132 for proteasomal inhibition for 5 hours before collection. While the DMSO and the Bafilomycin A1 treated groups still displayed lower endogenous 4EBP1 protein levels in the SIRT2 knockdown cells, the difference in 4EBP1 protein expression between control and SIRT2 knockdown cells was not observed in the MG132 treated group (Figure 4E). This result shows that SIRT2 affects the degradation of 4EBP1 through the proteasome. To further confirm this finding, ubiquitination of 4EBP1 was analyzed given that this post-translational modification leads to proteasomal degradation^53^. Flag-tagged 4EBP1 was transfected into control and SIRT2 knockdown HEK293T cells, and cells were treated with 40 μM MG132 for 6 hours before collection. After immunoprecipitating Flag-tagged 4EBP1, western blotting was used to analyze the ubiquitination level. As expected, 4EBP1 in SIRT2 knockdown HEK293T cells show a greater extent of ubiquitination compared to control cells, confirming that knocking down SIRT2 promotes the ubiquitination and degradation of 4EBP1 (Figure 4F).

### SIRT2 deacetylates 4EBP1 at K69 to stabilize 4EBP1

To further verify 4EPB1 as a SIRT2 substrate, we examined if SIRT2 interacts with 4EBP1. When co-expressed in HEK293T cells, HA-tagged SIRT2 was co-immunoprecipitated with Flag-tagged 4EBP1 (Figure 5A), suggesting that 4EBP1 interacts with SIRT2.

**Figure 5.**
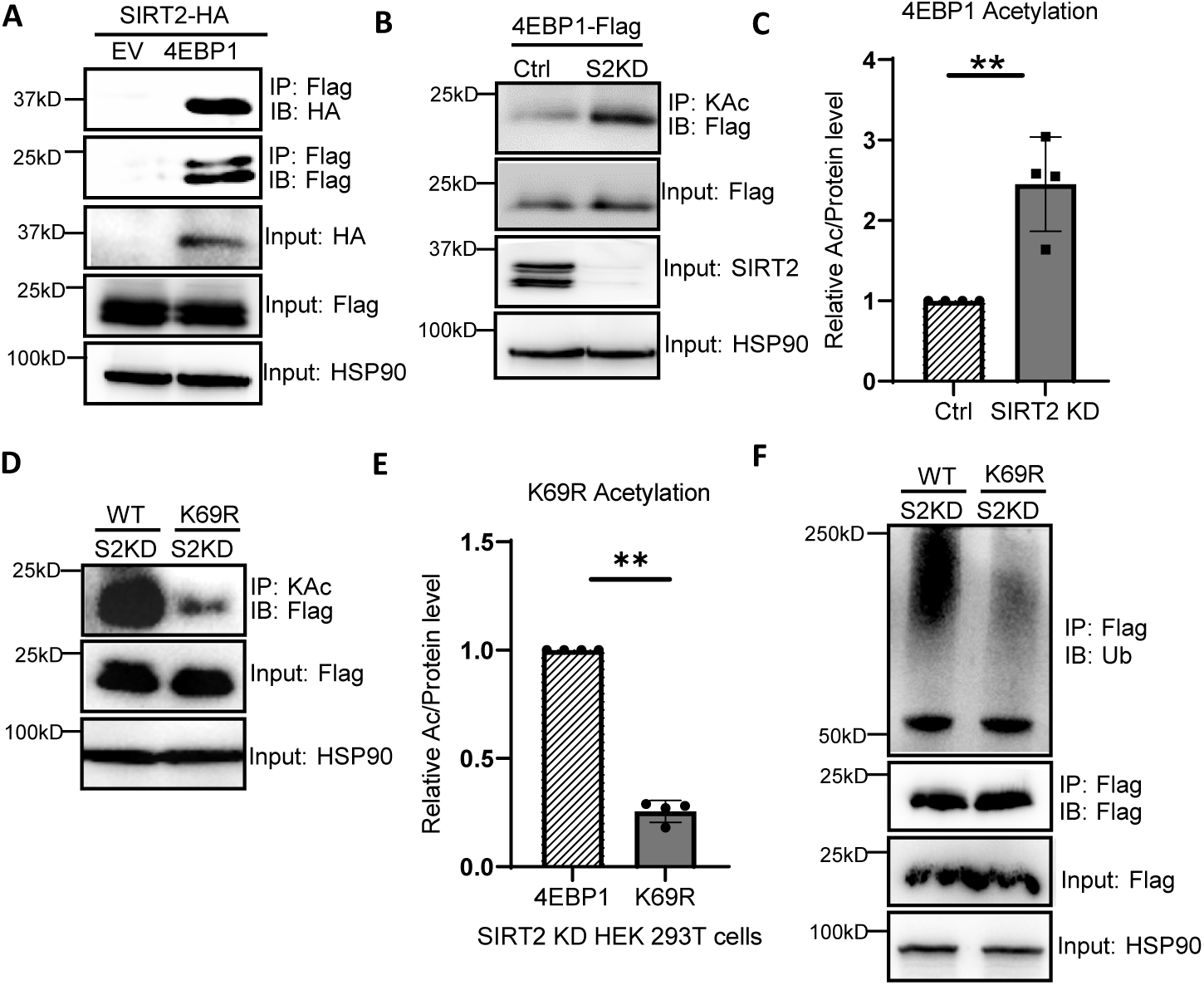
Deacetylation of 4EBP1 at K69 by SIRT2 stabilizes 4EBP1. **(A)** SIRT2 interacts with 4EBP1. HEK293T cells were transfected with HA-tagged SIRT2 along with Flag-tagged empty vector or Flag-tagged 4EBP1. Flag beads were used to pull down Flag-tagged proteins, and samples were analyzed using western blot. **(B)** 4EBP1 is deacetylated by SIRT2. Flag-tagged 4EBP1 was transfected into control and SIRT2 knockdown HEK293T cells. Cells were treated with 40 μM MG132 for 6 hours before collection. Acetylation was determined using acetyl lysine IP pull down and analyzing western blot for Flag-4EBP1. **(C)** Quantification of relative 4EBP1 acetylation levels between control and SIRT2 knockdown HEK293T cells. Error bars represent ±SD for experiments performed in quadruplet. ** indicating p<0.01. **(D)** SIRT2 deacetylates 4EBP1 at K69. Flag-tagged 4EBP1 or Flag-tagged 4EBP1 K69R mutant was transfected into SIRT2 knockdown HEK293T cells and pulled down with acetyl lysine IP beads. Cells were treated with 40 μM MG132 for 6 hours before collection. Acetylation was detected using western blot. **(E)** Ratios of 4EBP1 acetylation level to protein level were quantified and compared between 4EBP1 WT and K69R mutant in SIRT2 knockdown HEK293T cells. Error bars represent ±SD for experiments performed in quadruplet. ** indicating p<0.01. **(F)** Deacetylation of 4EBP1 at K69 by SIRT2 decreases ubiquitination. Flag-tagged 4EBP1 WT or K69R mutant was transfected into SIRT2 knockdown HEK293T cells. Cells were treated with 40 μM MG132 for 6 hours before collection. Ubiquitination level of 4EBP1 was analyzed by Flag IP pull down and western blot.

We then examined if 4EBP1 is acetylated and, if so, whether it can be deacetylated by SIRT2. Flag-tagged 4EBP1 was transfected into control and SIRT2 knockdown HEK293T cells. Since SIRT2 affects 4EBP1 proteasomal degradation, cells were treated with 40 μM of MG132 for 6 hours before collection to ensure the inhibition of degradation. Acetylated proteins were precipitated using anti-acetyl lysine beads, and then anti-Flag western blotting was performed to analyze the acetylation level of 4EBP1. Knocking down SIRT2 increased the acetylation level of 4EBP1 (Figure 5B). Quantification of four independent replicates showed that knocking down SIRT2 increases the acetylation of 4EBP1 by approximately 2.5-fold (Figure 5C), thus supporting that SIRT2 deacetylates 4EBP1.

After verifying that 4EBP1 is a substrate of SIRT2, we aimed to identify the deacetylate site of 4EBP1 by SIRT2, which could further help to elucidate the function of the deacetylation. 4EBP1 contains three lysine residues: K57, K69, and K105. Single, double, and triple lysine-to-arginine (K to R) mutants were prepared to determine the deacetylation site. The K69R mutant had a significantly decreased acetylation level compared to wild type 4EBP1 (Figure 5D and 5E), suggesting that K69 is the major site of deacetylation by SIRT2. Since we discovered that SIRT2 affects the ubiquitination of 4EBP1, we hypothesized that this effect could be through deacetylation at K69. To test this, Flag-tagged 4EBP1 or Flag-tagged 4EBP1 K69R mutant was transfected into SIRT2 knockdown HEK293T cells, and then the cells were treated with 40 μM of MG132 for 6 hours before collection. Flag-tagged proteins were immunoprecipitated using anti-Flag beads, and the ubiquitination levels were analyzed by western blotting. The K69R mutant showed a marked reduction in ubiquitination compared to the wildtype 4EBP1, indicating that deacetylation of 4EBP1 at K69 inhibits the ubiquitination of 4EBP1 (Figure 5F). This result also agrees with the observation that there was more endogenous 4EBP1 in control cells than in SIRT2 knockdown cells. Taken together, our data suggest that SIRT2 deacetylates 4EBP1 at K69, and this deacetylation inhibits 4EBP1 ubiquitination, preventing it from being degraded through the proteasome, which in turn decreases protein translation (Figure 6).

**Figure 6.**
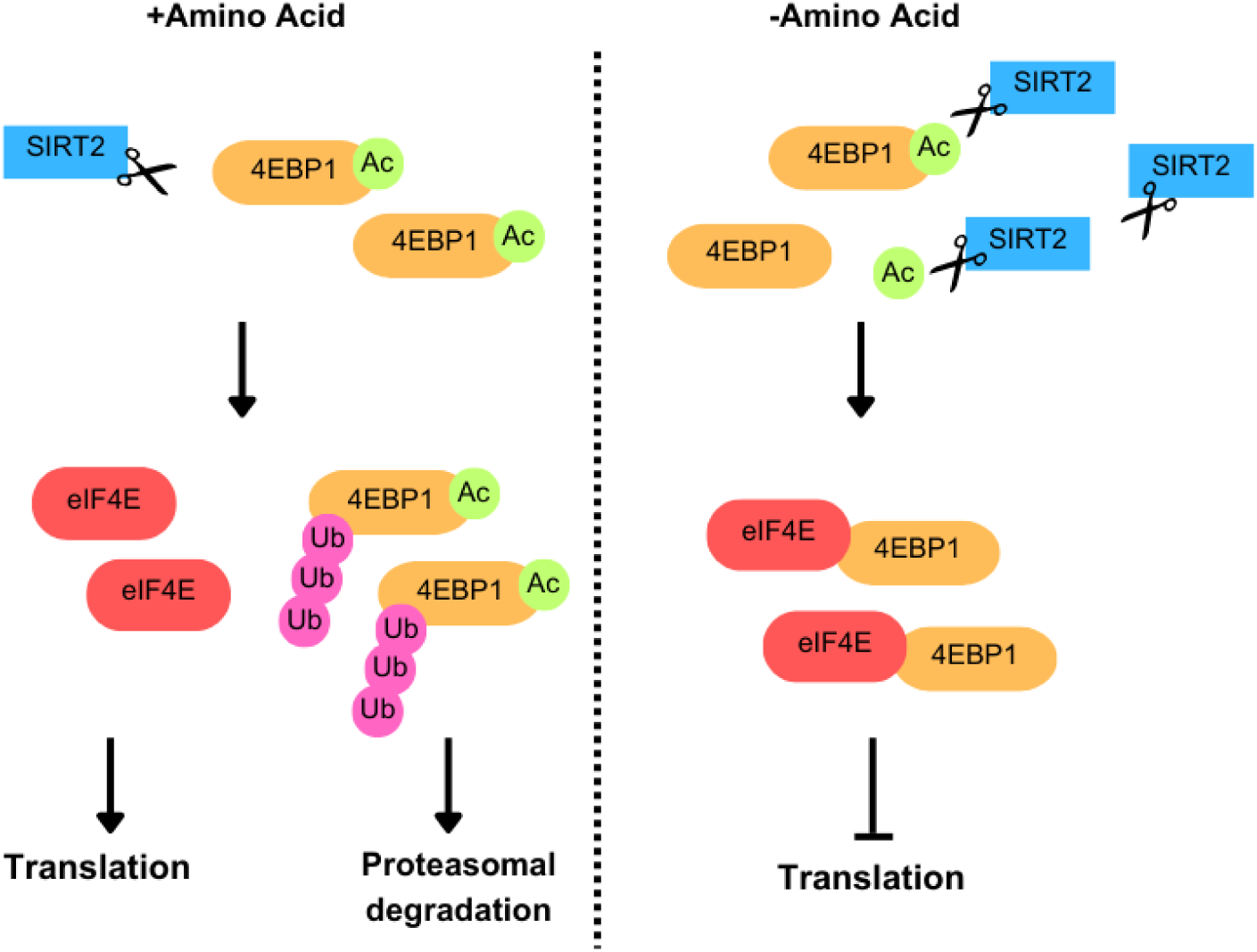
Working Model. Under amino acid limitation, SIRT2 protein level is upregulated, and 4EBP1 is more prone to deacetylation by SIRT2. SIRT2 deacetylates 4EBP1 at K69, which inhibits its ubiquitination and proteasomal degradation. The accumulated 4EBP1 binds to eIF4E, inhibiting protein translation. When amino acids are abundant, less SIRT2 is present, and more 4EBP1 is acetylated. Acetylated 4EBP1 is more ubiquitinated, which leads to proteasomal degradation. Because of the decreased endogenous 4EBP1, translation is upregulated.

## Discussion

Our study shows that SIRT2 is upregulated translationally under amino acid stress via its 5’uORF. The upregulated SIRT2 then suppresses translation by deacetylating and stabilizing the α form of 4EBP1. The results further broaden our understanding of SIRT2 as a stress response protein. SIRT2 is known to be important in responding to oxidative stress^54–56^, mitotic stress^57^, replication stress^58^, and Golgi stress^17^. It has been reported to be upregulated under oxidative stress and deacetylates the transcription factor FOXO3a, which in turns increases FOXO3a transactivation activity, reduces cellular ROS levels, and increases cell death under severe oxidative stress^54^. Although this study reported that SIRT2 levels are elevated under caloric restriction, the exact mechanism involved was not known. Here we provide a more detailed understanding of the upregulation of SIRT2 levels when cells are confronted with nutrient stress. We discovered that amino acid limitation triggers the elevation of SIRT2 levels in cells via the integrated stress response pathway. The upregulation of SIRT2 is similar to that of ATF4. Bioinformatics studies have shown that nearly half of the human transcriptome contain uORFs^41^. However, very few mRNAs have been reported to be directly regulated by uORF. Our study identified Sirt2 as a new mRNA transcript that is regulated by the uORF in response to amino acid limitations, thus expanding the knowledge of uORF-mediated translational regulation. We propose that like ATF4, SIRT2 is an important target upregulated by ISR. ATF4, as a transcription factor, helps to coordinate the stress response by controlling transcription. On the other hand, SIRT2, as an enzyme that modifies various substrate proteins, helps to coordinate the stress response at the post-translational level.

Suppress global protein translation under amino acid limitation serve to preserve energy and resources^42,59,60^. Therefore, we propose the model that SIRT2 is upregulated under amino acid limitation, which then suppresses global protein translation as a stress response to promote cell survival under amino acid limitation (Figure 6). Translation is subjected to complex regulation and crucial to cell survival and overall health. Our study sheds light on a key regulatory mechanism by which cells survive amino acid deprivation, by establishing 4EBP1 as a substrate for SIRT2 and the role SIRT2 plays in regulating protein translation.

4EBP1 has long been known as an important player in protein translation, and its phosphorylation has been widely studied. The phosphorylation of 4EBP1 by mTORC1 causes 4EBP1 to dissociate from eIF4E, which then promotes translation ^12,13^. However, the regulation of 4EBP1 by acetylation has not been previously reported. We found that 4EBP1 is acetylated on K69, and that this acetylation was removed by SIRT2. K69 acetylation promotes 4EBP1 ubiquitination and proteasomal degradation, whereas, SIRT2, by deacetylating K69, stabilizes 4EBP1 and thus inhibits translation. Ubiquitination of 4EBP1 is reported to occur on K57, but how the ubiquitination level is regulated has not been understood ^61^. Our work shows that K69 acetylation decreases the ubiquitination level of 4EBP1. This finding fits into our model nicely. When the SIRT2 protein level is upregulated under conditions of amino acid deprivation, 4EBP1 will be more likely to be deacetylated by SIRT2 at K69 and become less ubiquitinated at K57, thus stabilizing its expression in cells. As a result, more endogenous 4EBP1 is available to bind to eIF4E, thus inhibiting translation.

While we focus on 4EBP1 as a SIRT2 substrate, which underlies the regulation of translation by SIRT2, we recognize that SIRT2 might suppress translation via other substrate proteins. Indeed, in another manuscript under joint consideration, it is found that SIRT2 suppresses translation via deacetylation and the inhibition of RheB, a small GTPase that is important for mTORC1 activation. This further supports the model that under ISR, SIRT2 helps to coordinate the stress response at the post-translational level by modifying different substrate proteins.

We note that in addition to oxidative stress and the integrated stress response mentioned above, SIRT2 is also upregulated by Golgi stress^17^. Thus, SIRT2 can be considered as an important stress response protein. With its levels/activities upregulated by various stresses, SIRT2 plays a critical role to help cells survive these stresses. Interestingly, SIRT2 inhibitors have been recognized to have broad anticancer activity without showing significant toxicity in normal cells^62^. The stress response role provides a logical explanation for the broad and cancer cell-selective toxicity of SIRT2 inhibitors, as cancer cells likely experience more stresses in order to proliferate and metastasize. Thus, our study provides a better understanding of the physiological function of SIRT2 and the utility of SIRT2 inhibitors in treating human diseases.

## Methods

### Reagents, Antibodies, and Plasmids

Anti-FLAG affinity gel (A2220), Thapsigargin, cycloheximide, and puromycin were purchased from Sigma-Aldrich. Dual-Luciferase® Reporter Assay System was purchased from Promega. TM is synthesized as previously described^62^. Polyethyleneimine (PEI, 24765) was purchased from Polysciences. Acetyl-Lysine Affinity Beads (AAC04) was purchased from Cytoskeleton, Inc. MG132 and Bafilomycin A1 were purchased from MedChemExpress.

The following antibodies were purchased from Cell Signaling Technology: SIRT2 (D4050), SIRT1 (D739), SIRT3 (D22A3), SIRT5 (D8C3), SIRT6 (D8D12), HSP90 (C45G5), ATF4 (D4B8), p-S6K Thr389 (9205), S6K (9202), p-4EBP1 Ser65 (9451), 4EBP1 (9452), Ubiquitin (E4I2J), HA-Tag (C29F4), Anti-rabbit/mouse IgG HRP-linked Antibody. The following antibodies were from Santa Cruz: SIRT7 (C-3), β-actin (C-4). The following antibodies were from Sigma: anti-Flag M2 antibody conjugated with horseradish peroxidase (A8492), anti-puromycin (12D10).

Luciferase constructs were purchased from Promega. The human 302 bp *Sirt2* 5’UTR was amplified by PCR from cDNA library and inserted between HindIII and NcoI in pGL3 vector using primers 5’-AGT CAG <u>AAG CTT</u> CAT TTT CCG GGC GCC -3’ and 5’-AGT CAG <u>CCA TGG</u> GCG CGG -3’. The start codon of *SIRT2* main ORF was fused in frame with *firefly*. To disrupt uORF (133-234), site directed mutagenesis was carried out using 5’-GTC TGC GGC CGC AAT <u>A</u>TC TGC TGA GAG TTG T -3’ and 5’-<u>T</u>AT TGC GGC CGC AGA CGC GCT TTC GTA CAA C -3’.

Flag tagged 4EBP1 wildtype and KR mutant vectors were purchased from GenScript. GenScript cloned 4EBP1 gene into pcDNA3.1+/C-(K)-DYK vector and then performed mutagenesis to generate the KR mutants. SIRT2 wildtype expression vector with FLAG tag was made as previously described^63^.

### Immunoblotting

Plasmids encoding Flag or HA tagged proteins were transfected into 293T control and SIRT2 knockdown cells using PEI. For the transfection process, 6 μg of plasmids were first added to 1 mL of DMEM media, then 18 μl of PEI was added to the mixture. The mixture was incubated for 15 minutes at room temperature before added to the cells. An empty plasmid sometimes was used as a negative control. Cells were collected the day after the transfection and were washed with PBS prior to collecting via centrifugation of 3000 rpm for 5 minutes at 4 °C. For each sample, 1 ml of 1% NP40 lysis buffer with final pH 7.4 (150 mM Tris-HCl, pH 8, 150 mM NaCl, 10% glycerol, 1% NP40) was used to lyse the cells at 4 °C for 1 hour. 100 μl lysis buffer was used for 6 well plate samples. After spinning down the samples with 13000 rpm for 10 minutes at 4 °C, the supernatant was collected and normalized after using the Bradford reagent to determine the protein concentration. As input, 40 μl of each normalized sample were collected, mixed with 8 μL of 6x loading dye, boiled at 95 °C for 5 minutes. For checking endogenous protein, the samples were then proceeded to standard western blot procedure. For the immunoprecipitation experiments, the rest of the normalized sample were incubated with 20 μl of HA, Flag, or acetyl lysine beads at 4 °C overnight. The affinity beads were then washed with IP washing buffer (25 mM Tris, pH 8.0, 150 mM NaCl, 0.2% NP40) for 3 times and were dried with a 1 ml syringe. Depending on the experiment, 50 μl of 1x loading dye or 35 μl of 2x loading dye were added to the dried beads, and each sample was boiled at 95 °C for 5 minutes. Standard western blot analysis was used to analyze the samples. Western blots were performed as previously described^64^. The proteins were detected using enzyme linked fluorescence (ECL, pierce biotechnology Inc., Clarity Max, Bio-Rad; Immobilon Western Chemiluminescent HRP Substrate, Millipore) on a ChemiDoc (Bio-Rad).

### Cell Culture

HEK293T, MCF7, Hela, and HAP1 cells were grown in DMEM media (Invitrogen) supplemented with 10% (vol/vol) fetal bovine serum (FBS; Invitrogen, Carlsbad, CA). MEF cells were cultured in DMEM supplemented with non-essential amino acids (Invitrogen) and 15% FBS. A549 cells were cultured in RPMI-1640 (Invitrogen) with 10% FBS. HCT116 cells were cultured in McCoy’s 5A with 10% FBS. For amino acids depletion experiments, DMEM, RPMI, and McCoy’s 5A depleted of indicated amino acids supplemented with the indicated percent of dialyzed FBS (Invitrogen), while the control groups were cultured in full media also with dialyzed FBS.

SIRT2 control and stable knockout MEF cells, SIRT2 control and stable knockdown HEK293T cells were generated as previously described^64^.

### RT-PCR analysis of mRNA levels

Total mRNA was extracted using RNeasy Mini Kit (Qiagen, CA, USA) according to the manufacturer’s instructions and then reverse transcribed to cDNA library using SuperScript Vilo cDNA Synthesis Kit (Thermo Fisher). Real-time PCR were performed on QuantStudio™ 7 Flex Real-Time PCR System using SYBR™ Green PCR Master Mix (Applied Biosystems) and primers for *SIRT2*: 5’-TGC GGA ACT TAT TCT CCC AGA -3’, 5’-GAG AGC GAA AGT CGG GGA T - 3’; *ATF4*: 5’-GTT CTC CAG CGA CAA GGC TA -3’, 5’-ATC CTG CTT GCT GTT GTT GG -3’; *ACTB* (internal control): 5’-CAT GTA CGT TGC TAT CCA GGC -3’, 5’-CTC CTT AAT GTC ACG CAC GAT -3’.

### Luciferase Assay

Firefly and renilla constructs were co-transfected at 10:1 ratio using FuGene 6 into Hela cells in full DMEM media. After 12 hrs the cells were exposed to indicated stress and then collected and analyzed using Dual-Luciferase Reporter Assay System according to manufacturer’s instructions.

### Data Analysis

Prism (Graphpad Software) was used for data analysis. The two-tailed Student’s t test was used to determine statistical significance between two groups. Experimental values are shown as mean ±SD. *p<0.05, ** p<0.01, *** p<0.001.

## Author contributions

Y.Z.: designed and analyzed the experiments as well wrote and edited the manuscript. M.W: designed and analyzed the experiments. D.H.: designed and analyzed experiments involving SIRT2 inhibition or overexpression and translation. R.C.: conceptualization and manuscript editing. H.L.: conceptualization and manuscript editing.

## Lead Contact

Additional information and requests for materials should be directed to the lead contact Hening Lin.

---

**Figure S1.**
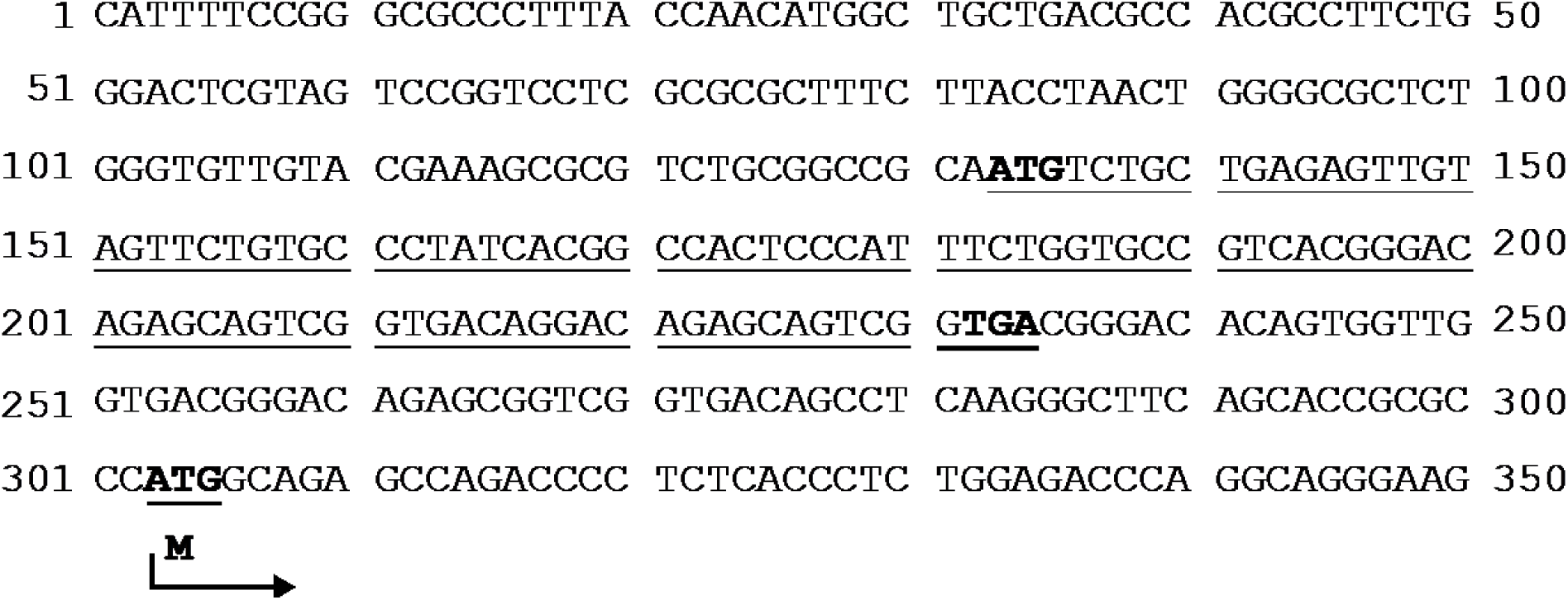
*5’UTR from the human Sirt2 variant 1*.

**Figure S2.**
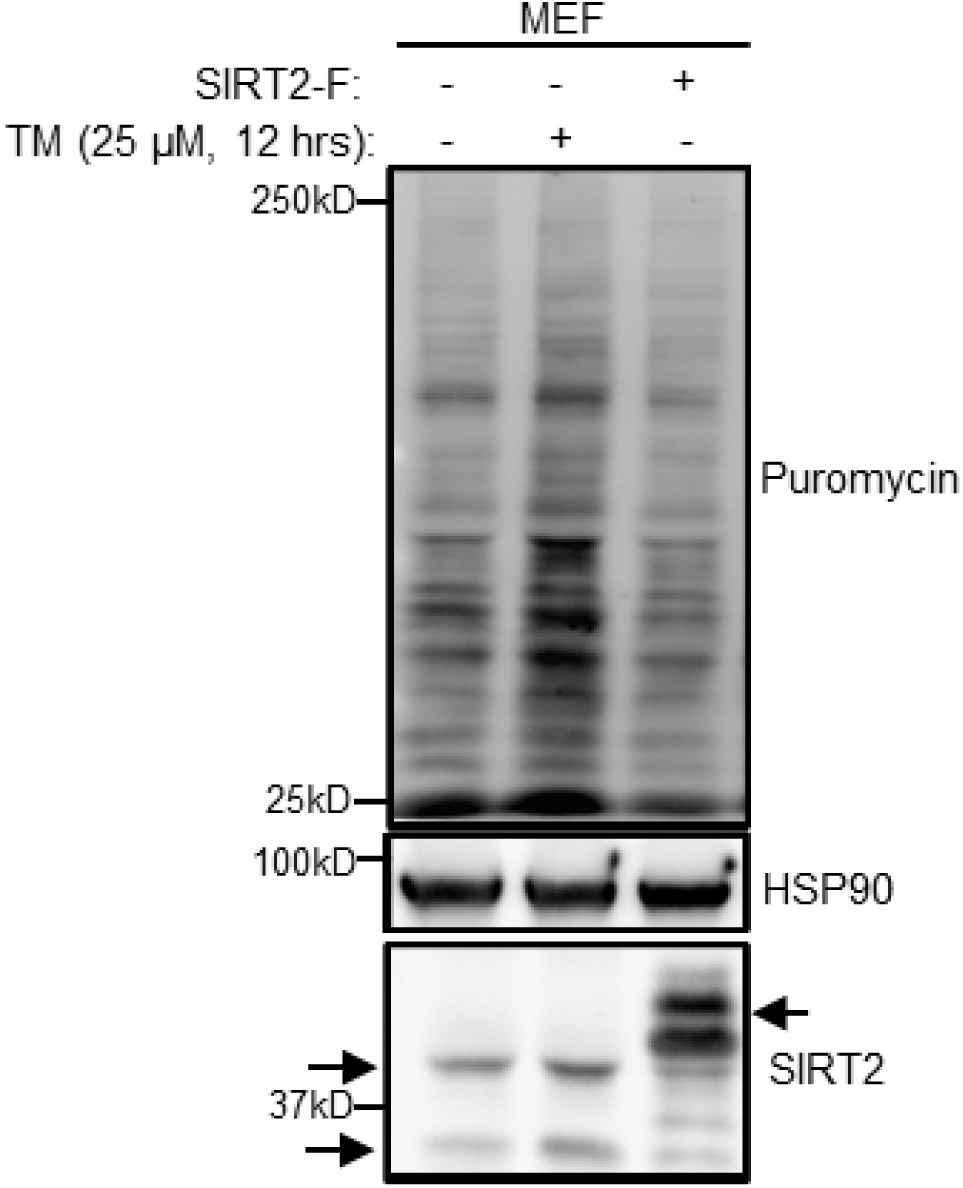
SIRT2 inhibition promotes protein translation, and SIRT2 overexpression downregulates protein translation. MEF cells were either treated with 25 μM of SIRT2 inhibitor TM for 12 hours or overexpressed with Flag-tagged SIRT2. Cells were treated with 10 μg/ml puromycin for 10 minutes before collecting. Results were analyzed using western blot.

**Figure S3.**
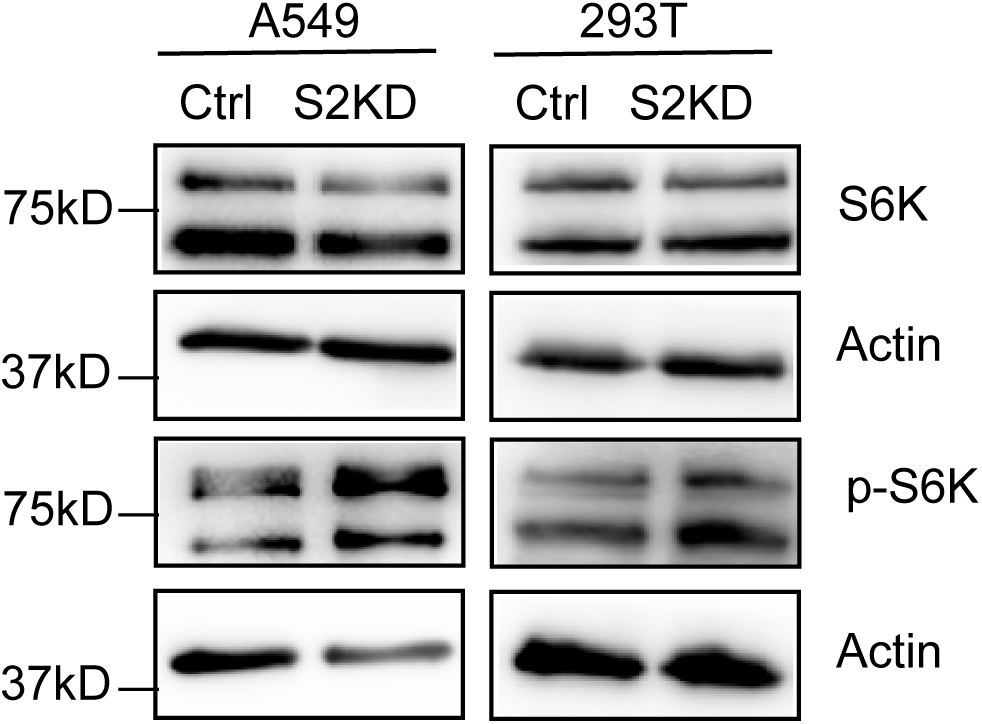
S6K and p-S6K levels in control and SIRT2 knockdown A549 and HEK293T cells. Protein levels were measured from whole cell lysate using western blot.

**Figure S4.**
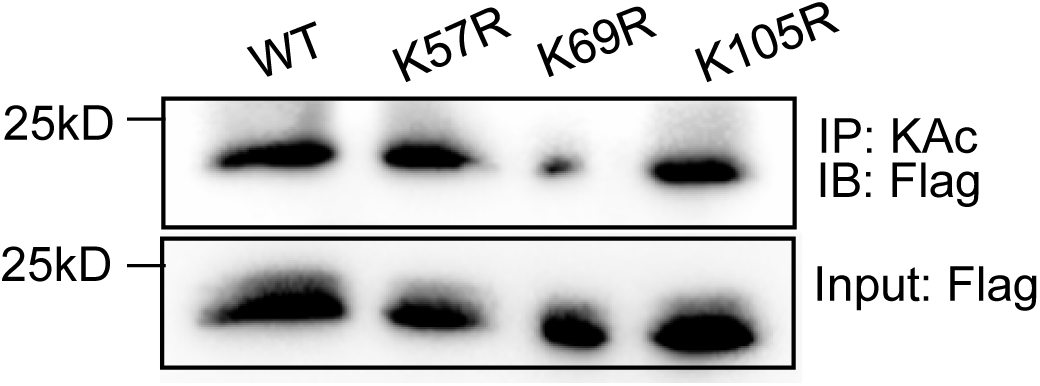
SIRT2 deacetylates 4EBP1 only at K69. SIRT2 knockdown HEK293T cells were transfected with Flag-tagged 4EBP1 WT, K57R, K69R, or K105R mutant. Acetylated proteins were pulled down with acetyl lysine IP beads. Acetylation was detected using western blot.

**Figure S5.**
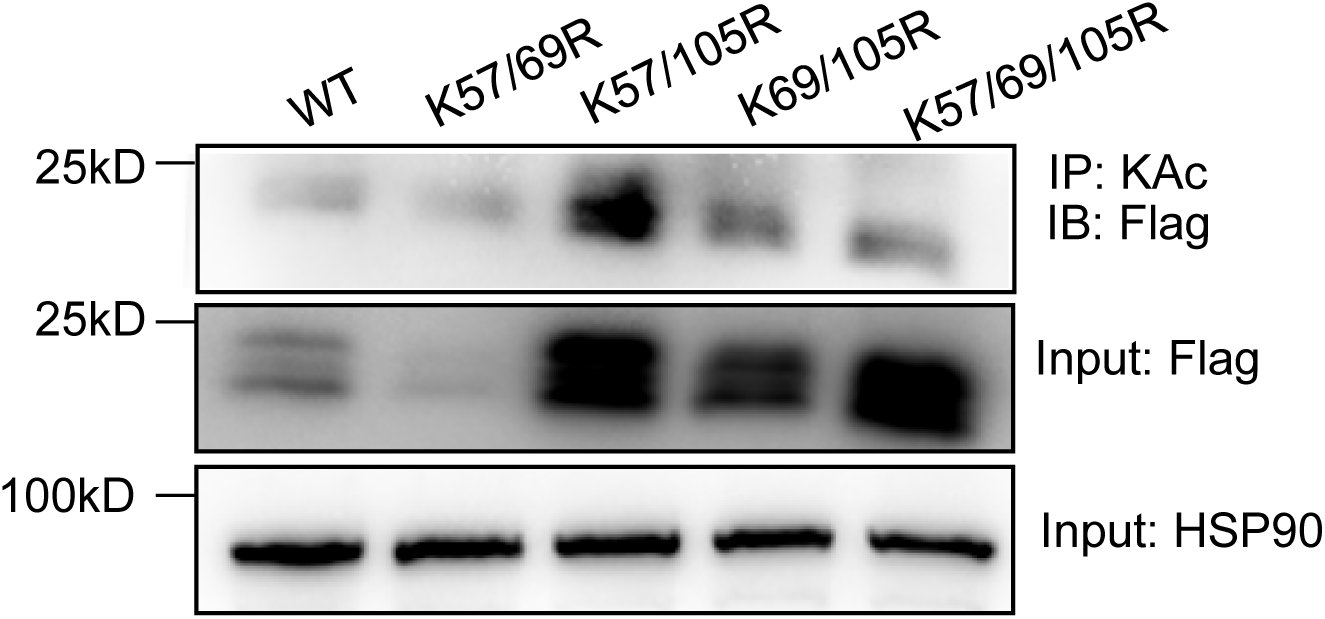
SIRT2 deacetylates 4EBP1 only at K69. SIRT2 knockdown HEK293T cells were transfected with Flag-tagged 4EBP1 WT, double, or triple mutants (K57/69R, K57/105R, K69/105R, K57/69/105R). Acetylated proteins were pulled down with acetyl lysine IP beads. Acetylation was detected using western blot.

